# Genome-wide aggregated *trans*-effects analysis identifies key role in hypertension for proteins involved in clearance of triglyceride-rich lipoproteins

**DOI:** 10.64898/2026.09.04.749549

**Authors:** Svitlana Braichenko, Andrii Iakovliev, Jay Dyer, Xuan Zhou, Athina Spiliopoulou, Helen M Colhoun, Paul M McKeigue

## Abstract

**Background:** Genome-wide aggregated *trans*-effects (GATE) analysis is a novel method in which *trans*-effects on gene expression (as transcript or protein) are combined with SNP-trait association data to identify core genes that directly influence the trait. The objective of this study was to identify core genes for blood pressure.

**Methods:** We undertook GWASs of mean arterial pressure (MAP) and and body mass index (BMI) in 373,882 individuals aged less than 60 in the Our Future Health (OFH) cohort. Using summary statistics from GWASs of circulating proteins on the SomaScan and Olink platforms, we tested for association of GATE scores (predicted levels of each protein based on *trans*-effects) with MAP. We confirmed replication of top associations in an independent cohort.

**Results:** The strongest GATE score association with MAP was for *LPL* (lipoprotein lipase). Higher genetically predicted circulating levels of *LPL* were associated with lower MAP but higher BMI. GATE scores for three other proteins involved in lipid handling – *CD300LG*, *ADIPOQ*, *TIMP4* – were also associated with lower MAP but higher BMI. GATE scores for all four of these proteins were inversely associated with chylomicron triglyceride (measured as XXL-VLDL-TG by NMR spectroscopy). The associations of GATE scores for *ANGPTL3* and *ANGPTL4*, which regulate *LPL*, were consistent with a causal role of *LPL* in lowering XXL-VLDL-TG and MAP.

**Conclusions:** These results point to a key role in hypertension for proteins that regulate post-prandial clearance of triglyceride-rich lipoproteins, independently of adiposity.

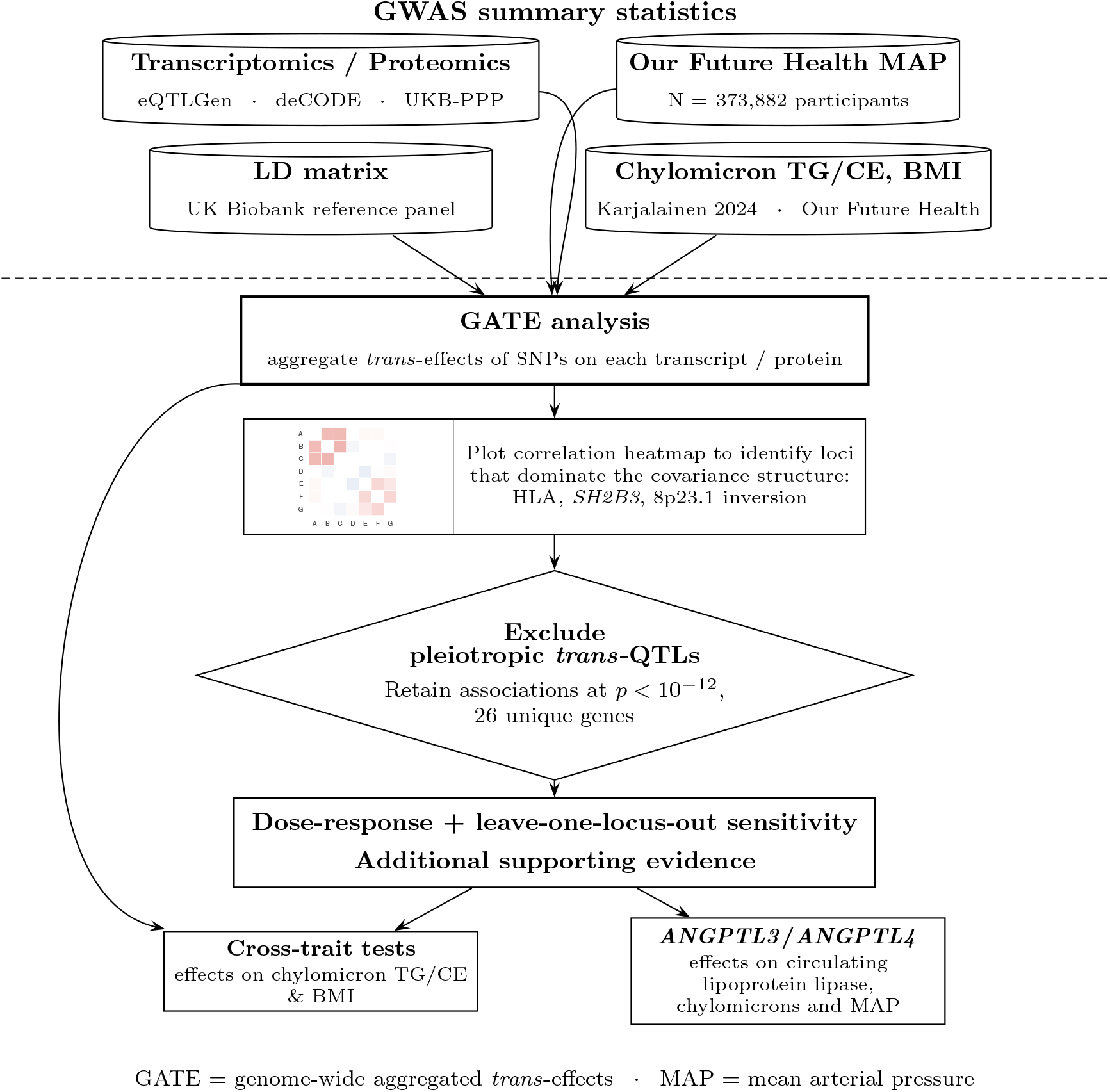

## Introduction

Genome-wide association studies (GWAS) have identified more than 1000 regions in which common variants influence blood pressure traits ^1^. However going from GWAS discovery to drug target identification has been challenging; most of the variants associated with blood pressure are non-coding and have small effect sizes. The nearby genes assigned as the most likely mediators of the effects of these variants on blood pressure often have little obvious relevance to blood pressure and few have led to drug development programmes. The “omnigenic” sparse effector hypothesis postulates that the polygenic effects of common SNPs on a typical complex trait coalesce via *trans*-effects on the expression of a sparse set of “core” genes that directly influence the trait ^2^. The availability of summary statistics from large genome-wide association studies of gene expression in whole blood or proteins in serum or plasma has made it possible to apply this model to identify core genes by genome-wide aggregated *trans*-effects (GATE) analysis. In this approach, summary statistics are used to learn weights for predicting levels of each transcript or protein from genotypes at quantitative trait loci (QTLs) that are distant from the site of the gene encoding the transcript or protein. In a target cohort, the GATE score for each transcript or protein in each individual is computed as the weighted sum of the individual’s genotypes. Each GATE score is then tested for association with the disease or trait under study ^3,4^. The objective of this study was to identify core genes for blood pressure using GATE analysis.

## Methods

### Source cohorts for blood pressure analyses

The discovery cohort in this study is Our Future Health (OFH), a study of more than 2 million adults in the UK of whom 755,000 have been genotyped with imputation against UK Biobank whole-genome sequence data as reference panel. We restricted the analysis to participants aged less than 60 years, in whom genetic effects on blood pressure are stronger and age-related comorbidities are less likely to confound associations ^5^. As summary statistics from large transcriptomic and proteomic studies are available only for populations of European ancestry, we restricted the analysis further to the 373,882 individuals of European ancestry. We calculated mean arterial pressure (MAP) as two-thirds diastolic plus one-third systolic blood pressure, excluding the first of three measurements at the assessment visit. For participants on antihypertensive therapy, untreated blood pressure was imputed by adding 15 mmHg to systolic and 10 mmHg to diastolic blood pressure ^6^. We undertook a GWAS using the REGENIE package restricting to SNPs with minor allele frequency greater than 1%, with age at assessment, sex and the first 20 genetic principal components as covariates. For replication we used summary statistics from a GWAS of systolic blood pressure in 99,785 individuals from the Genetic Epidemiology Research on Adult Health and Aging (GERA) cohort ^7^.

### Source cohorts for cross-trait analyses

Comparison of effects of GATE scores on blood pressure with effects on related traits such as obesity and post-prandial lipids can yield mechanistic insight. To study effects on body mass index (BMI) we undertook a GWAS of BMI in OFH participants aged under 60 years. To study effects on post-prandial lipids, we used summary statistics from a GWAS of “triglyceride levels in chylomicrons and extremely large very low density lipoprotein particles” (XXL-VLDL-TG) measured by NMR spectroscopy in 135,916 individuals (88% European ancestry) from 32 cohorts of which 26 were classified as fasted ^8^. We computed GATE scores for XXL-VLDL-TG and also for XXL-VLDL-CE (cholesteryl esters in XXL-VLDL) as that measurement was included in a published study of associations of with blood pressure ^9^

### Defining QTLs

For conciseness we use the term “exposure” for the transcript or protein measured on a specific platform and “outcome” for the trait under study. We extracted summary statistics for SNP associations with exposures, filtered to exclude SNPs not included in the outcome summary statistics, from three GWASs of transcriptomics or proteomics that were each based on more than 30,000 individuals: the eQTLGen Consortium Phase 1 meta-analysis of whole blood transcripts ^10^ in which only 10,317 trait-associated SNPs were tested for *trans*-effects; the deCODE proteomics study ^11^ (4886 proteins on the SomaScan platform); and the UK Biobank Pharma Proteomics Project (UKB-PPP) ^12^ (2884 proteins on the Olink platform). For each genomic region in which at least one SNP is associated with the exposure at *p* < 10^−6^, a QTL is defined by grouping all SNPs associated with that exposure at *p* < 10^−5^.

### GATE analysis

Methods for GATE analysis with individual-level outcome data have been described previously ^3,4^. In brief, the GATE score for an exposure is the genetically predicted level of the exposure in that individual, obtained by aggregating the effects of exposure-associated SNPs at all *trans*-QTLs. The GATE coefficient for the exposure is the coefficient of regression of the outcome on the GATE score. For this study we used summary statistics for SNP-outcome associations, rather than individual-level genotype and phenotype data; this allows us to compute leave-one-locus-out (LOLO) sensitivity analyses efficiently. A technical description of the method is given in the Supplementary Material.

For each GATE score association with MAP, we tested for a dose-response relationship between the QTL-exposure coefficients and the QTL-outcome coefficients as described previously ^13^. This is equivalent to a 2-step Mendelian randomization (MR) analysis using a Bayesian method to model the distribution of direct (pleiotropic) effects on QTLs on the outcome ^14^, with the difference that instead of reporting the Bayesian posterior mean we report the maximum likelihood estimate of the causal effect parameter and the *p*-value. As the level of a transcript or protein in blood does not necessarily correspond to the dose in trait-relevant tissues, the absence of a dose-response relationship does not preclude causality.

### Restricting to associations that are driven by many *trans*-QTLs

Association of a GATE score with an outcome is more likely to be causal if the GATE score is based on aggregating many *trans*-QTLs than if the score is dominated by a few *trans*-QTLs of large effect. An intuitive way to explain this is that if the exposure is causal the ratio of “signal” (causal effects that are consistent in direction) to “noise” (pleiotropic effects that are random in direction) should increase with the number of *trans*-QTLs. To restrict to associations that are driven by many *trans*-QTLs, we used three methods: (1) restriction of GATE scores by effective number of *trans*-QTLs; (2) LOLO sensitivity analyses to identify clusters of GATE scores for which the association with the outcome is dominated by a single *trans*-pQTL region of large effect; (3) exclusion of pleiotropic genomic regions that have *trans*-effects on multiple exposures and large effects on outcome.

We calculated for each exposure the effective number *N*_eff_ of unlinked *trans*-QTLs as 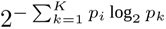 , where 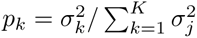 and *σ_k_* denotes the variance of the *k*th locus-specific *trans* score. This index ranges from 1 (when one QTL substantially dominates variance) to *K* (when locus-specific variances are approximately equal) ^15^. We restricted the analysis of *trans*-effects to the 3546 exposures with *N*_eff_*_>_*_5_, which cover 2895 distinct genes.

For each GATE score associated with the outcome, we recomputed the GATE coefficient *K* times, each time omitting one of the *K trans*-QTLs. We report the range of these LOLO estimates, the position of the most influential *trans*-QTL (the QTL for which omission produces the largest change in the GATE coefficient), the most likely gene mediating *in cis* the *trans*-effects of this QTL on the exposure, and the *p*-value for association with the residual GATE score when the most influential *trans*-QTL is omitted.

As in previous studies, we excluded *trans*-QTLs in the human leukocyte antigen (HLA) region (25 to 34 Mb on chromosome 6), which is a *trans*-QTL hotspot and has effects on multiple traits. In an initial analysis, we used two iterations of LOLO analyses to identify and exclude additional “peripheral master regulators”: genomic regions that dominate GATE associations for multiple exposures.

## Results

### GATE score associations with MAP

Two initial iterations of leave-one-locus-out (LOLO) analysis showed that GATE associations with MAP were dominated by *trans*-QTLs in the *SH2B3* region (108 to 113 Mb on chromosome 12) and the 8p23.1 genomic inversion region (8.23 to 12.04 Mb on chromosome 8). *Trans*-QTLs in these two regions were excluded from subsequent analyses for the reasons described above.

There were 61 exposures with *N*_eff_ *>* 5 for which GATE scores were associated with MAP at *p* < 10^−9^. We focus on the 27 associations with *N*_eff_ *>* 10 and *p* < 10^−12^ for association with MAP all of which are based on QTLs for proteins (pQTLs) rather than for transcripts (eQTLs). Table 1 shows these 27 GATE scores ordered by genomic position of the gene encoding the protein. For conciseness we use HUGO gene symbols to label proteins.

**Table 1.**
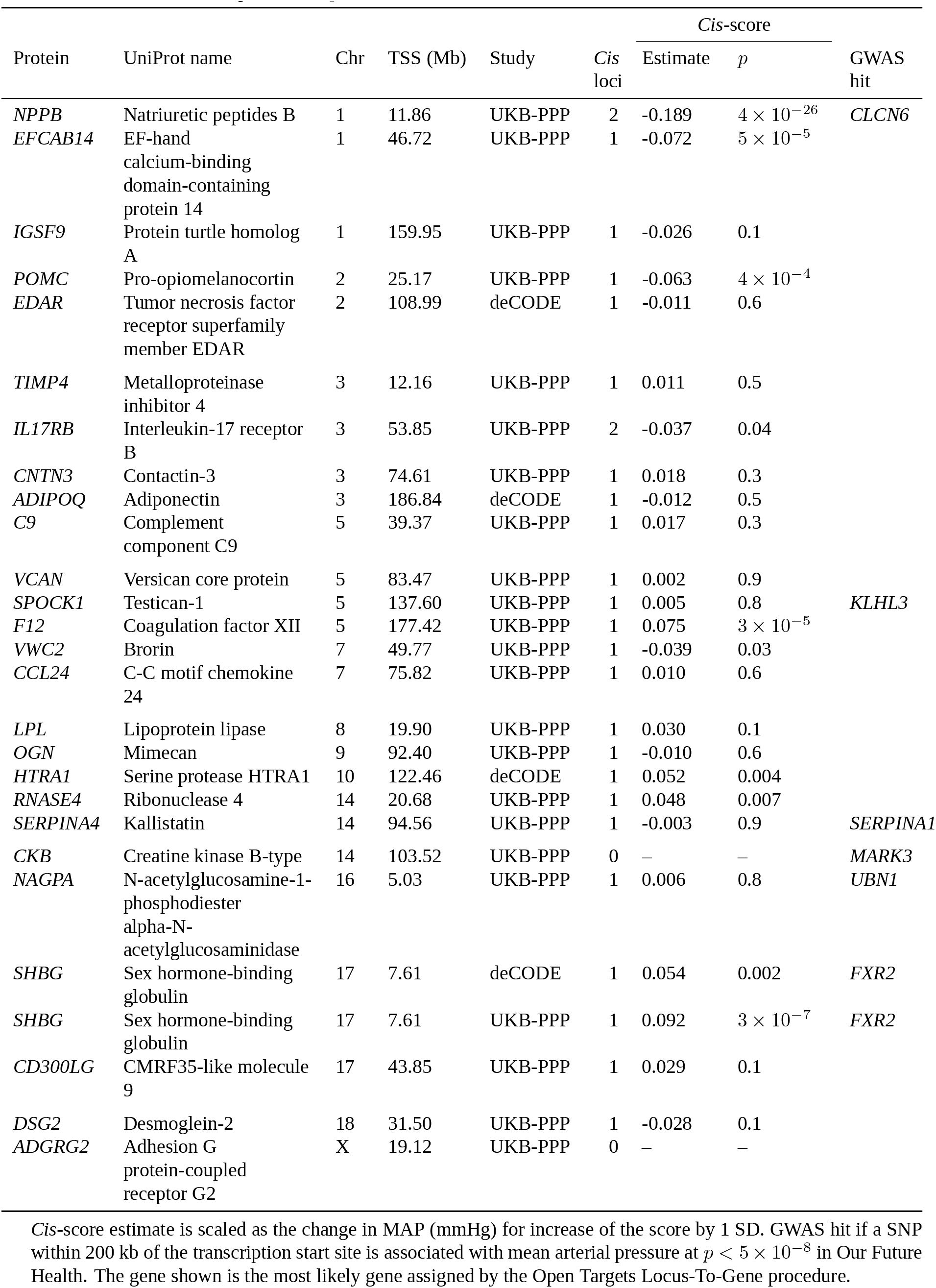
Trans-pQTL and cis-pQTLs for the 27 exposures with *N*_eff_ *>* 10 trans-pQTLs and GATE score associated with mean arterial pressure at *p* < 1 × 10^−12^.

Figure 1 shows a heat map of correlations between the GATE scores in Table 1, ordered by hierarchical clustering on the squared correlations so that correlated variables appear in blocks. Correlations between GATE scores indicate that the proteins share the same *trans*-pQTLs: in other words that they are genetically co-regulated. At the top of the correlation plot are six GATE scores that are almost uncorrelated: these include *POMC* (pro-opiomelanocortin), *NPPB* (natriuretic peptide B), *SPOCK1* (testican), and *VCAN* (versican). Below this is a block of weakly correlated scores for lipid-related genes including *ADIPOQ* (adiponectin), *LPL* (lipoprotein lipase), *CD300LG* (nepmucin) and *TIMP4* (metalloproteinase inhibitor 4). Below this are two more clusters, denoted for conciseness as the *F12* (coagulation factor XII) and *OGN* (osteoglycin) clusters.

**Fig 1.**
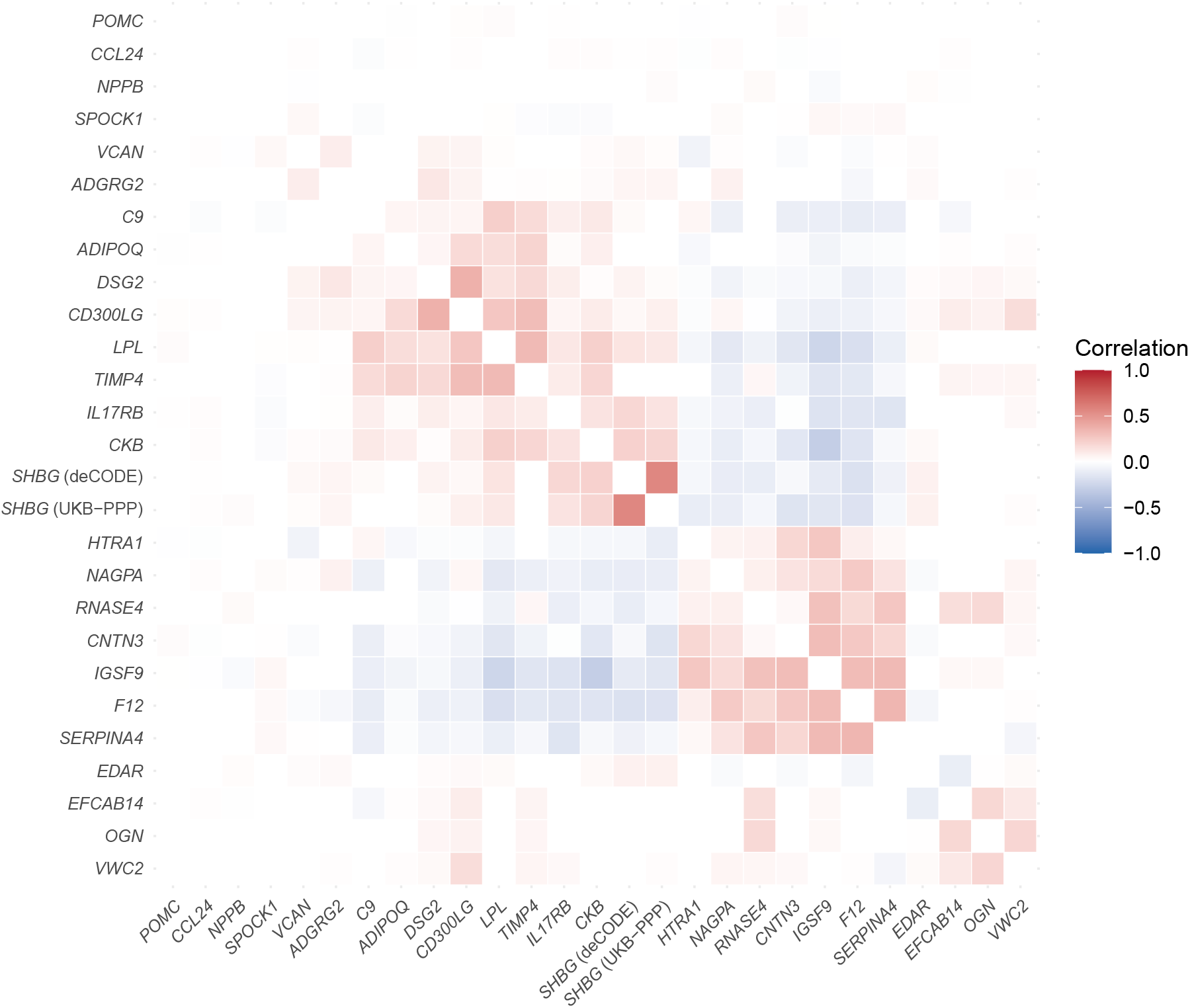
Correlation between GATE scores for the 27 exposures in Table 1 computed using the LD matrix between SNPs in UK Biobank participants of European ancestry. The *SH2B3* and 8p23.1 genomic inversion regions are excluded from the calculation of GATE scores. Rows and columns are ordered by hierarchical clustering on the squared correlations so that co-regulated genes appear in blocks.

Table 2 shows the GATE coefficients for these top 27 proteins, the LOLO range, and the dose-response analysis. Supplementary Table S1 shows that with the exception of *CCL24*, GATE scores for these proteins remained associated with MAP at *p* < 10^−5^ after leaving out the most influential locus. Supplementary Table S2 shows that associations with blood pressure of GATE scores for most of the proteins in the block of lipid-related genes are replicated in the GERA cohort.

**Table 2.** GATE and dose-response associations with MAP for the 27 exposures in Table 1 (*cis*-score associations are shown in columns 7-8 of that table).

| Protein | Study | *Trans* loci | $N_{\text{eff}}$ | GATE coefficient | | | MR dose-response | |
| --- | --- | --- | --- | --- | --- | --- | --- | --- |
| | | | | Estimate | $p$ | LOLO range | Estimate | $p$ |
| *POMC* | UKB-PPP | 26 | 12.1 | -0.163 | $9 \times 10^{-20}$ | [-0.188, -0.112] | -0.548 | 0.1 |
| *CCL24* | UKB-PPP | 33 | 19.7 | 0.154 | $7 \times 10^{-18}$ | [0.049, 0.169] | 0.225 | 0.2 |
| *NPPB* | UKB-PPP | 17 | 15.3 | -0.146 | $3 \times 10^{-16}$ | [-0.169, -0.113] | -1.036 | 0.02 |
| *SPOCK1* | UKB-PPP | 28 | 19.7 | -0.132 | $2 \times 10^{-13}$ | [-0.144, -0.113] | -0.516 | 0.02 |
| *VCAN* | UKB-PPP | 40 | 23.4 | -0.147 | $2 \times 10^{-16}$ | [-0.160, -0.085] | -0.325 | 0.04 |
| *ADGRG2* | UKB-PPP | 25 | 18.1 | -0.140 | $5 \times 10^{-15}$ | [-0.152, -0.109] | -0.438 | 0.1 |
| *C9* | UKB-PPP | 17 | 15.3 | -0.132 | $1 \times 10^{-13}$ | [-0.155, -0.105] | -0.646 | 0.2 |
| *ADIPOQ* | deCODE | 23 | 16.5 | -0.153 | $1 \times 10^{-17}$ | [-0.170, -0.113] | -0.575 | 0.02 |
| *DSG2* | UKB-PPP | 32 | 12.8 | -0.138 | $1 \times 10^{-14}$ | [-0.156, -0.103] | -0.342 | 0.2 |
| *CD300LG* | UKB-PPP | 43 | 30.0 | -0.137 | $2 \times 10^{-14}$ | [-0.153, -0.107] | -0.414 | 0.07 |
| *LPL* | UKB-PPP | 22 | 20.4 | -0.223 | $9 \times 10^{-36}$ | [-0.236, -0.206] | -1.777 | $6 \times 10^{-10}$ |
| *TIMP4* | UKB-PPP | 22 | 19.3 | -0.129 | $6 \times 10^{-13}$ | [-0.147, -0.110] | -0.787 | 0.09 |
| *IL17RB* | UKB-PPP | 34 | 29.5 | -0.166 | $2 \times 10^{-20}$ | [-0.172, -0.140] | -0.755 | $1 \times 10^{-4}$ |
| *CKB* | UKB-PPP | 22 | 19.4 | -0.133 | $1 \times 10^{-13}$ | [-0.165, -0.112] | -0.783 | 0.008 |
| *SHBG* | deCODE | 19 | 15.7 | -0.151 | $3 \times 10^{-17}$ | [-0.167, -0.101] | -0.619 | 0.08 |
| *SHBG* | UKB-PPP | 31 | 25.6 | -0.141 | $3 \times 10^{-15}$ | [-0.162, -0.100] | -0.493 | 0.07 |
| *HTRA1* | deCODE | 14 | 10.9 | 0.139 | $7 \times 10^{-15}$ | [0.097, 0.164] | 0.904 | 0.1 |
| *NAGPA* | UKB-PPP | 19 | 14.9 | 0.132 | $1 \times 10^{-13}$ | [0.110, 0.147] | 0.681 | 0.02 |
| *RNASE4* | UKB-PPP | 14 | 13.1 | 0.143 | $1 \times 10^{-15}$ | [0.115, 0.165] | 1.066 | 0.03 |
| *CNTN3* | UKB-PPP | 51 | 38.3 | 0.156 | $3 \times 10^{-18}$ | [0.135, 0.170] | 0.544 | $2 \times 10^{-4}$ |
| *IGSF9* | UKB-PPP | 16 | 10.7 | 0.133 | $1 \times 10^{-13}$ | [0.114, 0.145] | 0.972 | 0.004 |
| *F12* | UKB-PPP | 20 | 16.6 | 0.144 | $7 \times 10^{-16}$ | [0.109, 0.152] | 0.795 | 0.03 |
| *SERPINA4* | UKB-PPP | 15 | 12.5 | 0.128 | $8 \times 10^{-13}$ | [0.095, 0.139] | 0.793 | 0.03 |
| *EDAR* | deCODE | 20 | 13.0 | -0.130 | $3 \times 10^{-13}$ | [-0.140, -0.083] | -0.328 | 0.05 |
| *EFCAB14* | UKB-PPP | 13 | 12.3 | 0.167 | $8 \times 10^{-21}$ | [0.080, 0.189] | 0.851 | 0.3 |
| *OGN* | UKB-PPP | 24 | 21.6 | 0.154 | $9 \times 10^{-18}$ | [0.134, 0.174] | 1.108 | 0.009 |
| *VWC2* | UKB-PPP | 20 | 13.2 | 0.139 | $7 \times 10^{-15}$ | [0.120, 0.147] | 0.758 | 0.02 |

Table 1 shows for the genes encoding these top 27 proteins the associations of *cis*-scores with MAP. Where the transcription site of the gene is within 200 kb of SNPs associated with MAP at the conventional threshold of *p* < 5 × 10^−8^ in the OFH GWAS, the causal gene assigned by the Locus-To-Gene (L2G) procedure ^16^ (described in Supplementary Methods) for this GWAS hit is shown. For five of these genes there was a *cis*-score association with MAP at *p* < 0.001, and for six there was a GWAS hit within 200 kb. As *cis*-acting SNPs were excluded from the GATE scores, such *cis*-associations with MAP are supporting evidence of causality for genes identified through GATE score associations. The causal gene assigned by the L2G procedure did not agree with the gene encoding the GATE score protein for any of these genes. The strongest *cis*-score association was for *NPPB* (natriuretic peptide B), supported by a GWAS hit that had been attributed to *CLCN6*, 7 kb downstream of *NPPB*. This illustrates the uncertainty, for a highly polygenic trait such as blood pressure, in assigning the nearby causal gene that mediates a GWAS hit.

The strongest GATE score association with MAP was the inverse association with *LPL*, supported by a dose-response relationship between *trans*-pQTL effects on circulating lipoprotein lipase (LpL) and *trans*-pQTL effects on MAP, with an estimated causal effect size of -1.8 mmHg for an increase of one standard deviation (SD) in circulating lipoprotein lipase levels. Supplementary Figure S1 shows a scatter plot of the *trans*-pQTL coefficients underlying the dose-response model for LpL. The slope of the fitted line is the maximum likelihood estimate of the causal effect parameter. The statistical model allows for outliers to have direct effects on blood pressure that are not mediated through LpL.

For further illustration Figure 2 (bottom panel) shows the 22 *trans*-pQTLs that contribute to the GATE score for *LPL* and the 43 trans-pQTLs that contribute to the GATE score for *CD300LG*. Where a *trans*-pQTL is within 200 kb of a GWAS hit, the most likely gene assigned by the L2G procedure is labeled on the shared Manhattan plot at the top. Supplementary Table S3 shows for each of the pQTLs for *LPL* the nearby gene assigned as “most likely gene”. Of the 22 *trans*-pQTLs, in this table, 11 show inverse associations with MAP that are significant at *p* < 0.01. This coalescence on a target gene of multiple weak *trans*-effects that in aggregate are strongly associated with the outcome is the defining characteristic of a core gene. The corresponding pQTLs for *CD300LG* are shown in Supplementary Table S4.

**Fig 2.**
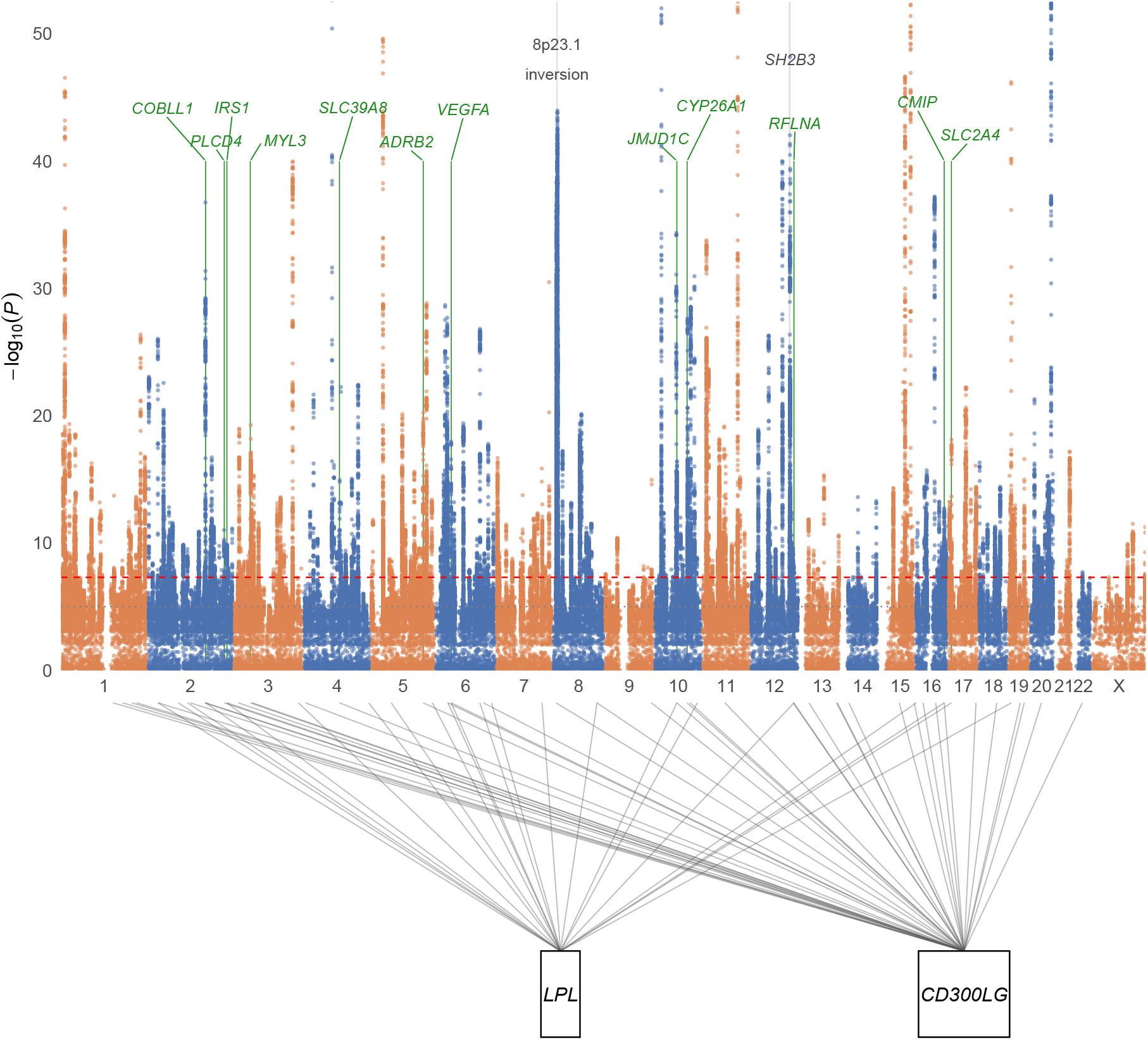
Top: Manhattan plot (truncated at − log_10_(*p*) = 50) of mean arterial pressure in 373,882 Our Future Health participants of European ancestry aged under 60 years; SNPs with − log_10_(*P* ) ≥ 4 kept in full, tapered random sample of the remainder shown for context. Thin green vertical lines : trans-pQTLs for *LPL* or *CD300LG* within 200 kb of a SNP association at *p* < 5 × 10^−8^), labeled with the most likely gene. Grey vertical lines mark the *SH2B3* and 8p23.1 inversion regions. Bottom: *trans*-pQTLs for *LPL* and *CD300LG*, each connected by a line from the genomic position of the pQTL to the position of the target gene.

### Cross-trait analyses: BMI and chylomicrons

As blood pressure is strongly related to obesity, we undertook a cross-trait analysis comparing, for the top 27 proteins identified through GATE effects on blood pressure, the GATE coefficients for blood pressure with the GATE coefficients for BMI. Figure 3A shows that for *LPL* and the other top three proteins in the lipid-related cluster – *CD300LG*, *ADIPOQ*, *TIMP4* – the *trans*-effects that raise the circulating levels of these proteins and lower blood pressure also raise body mass index. This indicates that their effects on blood pressure cannot be explained by shared genetic effects on adiposity.

**Fig 3.**
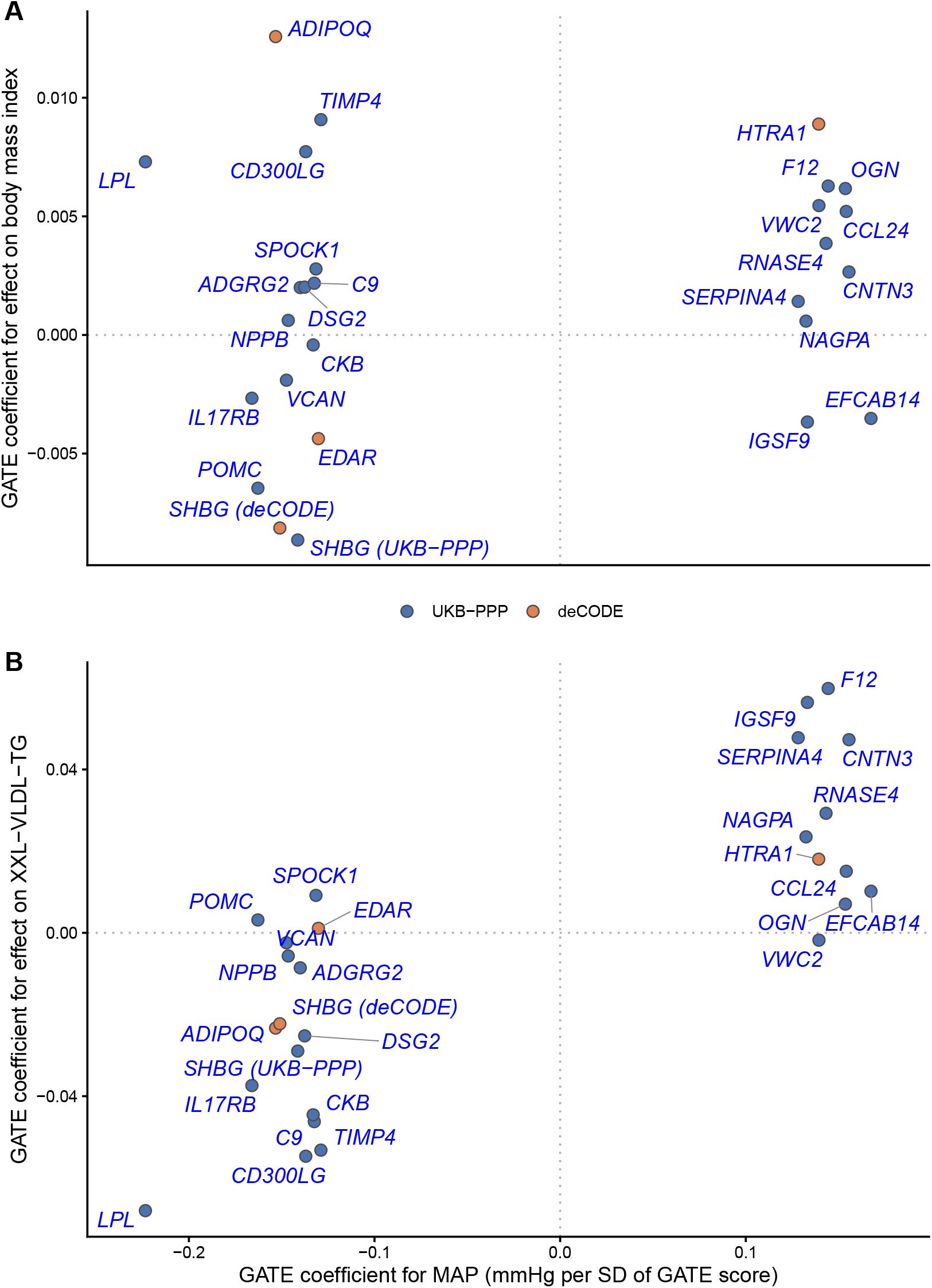
GATE coefficients for the 27 exposures in Table 2, effect on MAP (mmHg per SD of the GATE score) on the x axis (aligned between panels) against: (A) effect on body mass index (rank inverse-normal-transformed) in 373,519 Our Future Health participants of European ancestry aged under 60 years; (B) effect on triglyceride levels in chylomicrons and extremely large VLDL (XXL-VLDL-TG))(14).

As *LPL* has a key role in the clearance of post-prandial triglycerides from plasma, we next undertook a cross-trait analysis of XXL-VLDL-TG (a proxy for chylomicron triglyceride levels). We excluded pQTLs in the *APOA5* region (115,733 kb to 120,175 kb on chromosome 11) as initial analysis showed that GATE score associations with XXL-VLDL-TG were dominated by this region. Figure 3B shows that for most of the top 27 proteins the GATE coefficients for blood pressure and for XXL-VLDL-TG are concordant in direction. Crucially, the GATE coefficients for *LPL*, *CD300LG*, *ADIPOQ* and *TIMP4* are inversely associated with XXL-VLDL-TG, consistent with an effect on blood pressure mediated through effects on lipids, even though they are positively associated with body mass index. Supplementary Figure S2 shows that the GATE scores for XXL-VLDL-TG were highly correlated (*r* = 0.965) with GATE scores for XXL-VLDL-CE, for which associations with blood pressure have been reported ^9^.

### GATE score associations with blood pressure, body mass index and XXL-VLDL-TG

Supplementary Tables S6 and S7 show the GATE effects on MAP, XXL-VLDL-TG and BMI for a broader set of 60 exposures with *N*_eff_ *>* 5, GATE score associated with MAP at *p* < 10^−9^, and residual LOLO at *p* < 10^−5^. Of these 60 genes, 9 have support from a GWAS hit or rare variant association with blood pressure, and 7 have support from the effect of experimental pertubation of the gene or protein on blood pressure.

Supplementary Table S6. shows the GATE score associations for the 29 genes for which there is an effect on XXL-VLDL-TG that is concordant with the association with blood pressure. Except for *LPL*, *CD300LG*, *ADIPOQ* and *TIMP4*, the effects of most of these genes on MAP, BMI and XXL-VLDL-TG are concordant, suggesting that their effects are mediated at least partly through shared pathways regulating adiposity.

Supplementary TableS7 shows the GATE score associations for the 31 genes for which there is not an effect on XXL-VLDL-TG concordant with the effects on blood pressure. These include GATE scores for genes in the *OGN* cluster that have effects on BMI concordant with their effects on blood pressure: *POMC*, *HTRA1*, *EDAR*, *OGN* and *VWC2*. The coupling of genetic effects on these proteins with effects on blood pressure may thus be at least partly attributable to shared pathways that influence obesity.

For the remaining genes, which include *NPPB*, *SPOCK1* and *VCAN* the GATE scores have no effect on XXL-VLDL-TG or BMI. In this group we would expect to find genes that regulate blood pressure through pathways that do not depend on adiposity or lipid handling.

### GATE scores for angiopoietin-like proteins

LpL levels are regulated post-translationally by the angiopoietin-like proteins ANGPTL3, ANGPTL4 and ANGPTL8^17,18^. As ANGPTL3 and ANGPTL4 were measured in UKBB-PPP, we were able to study the relation of *trans*-effects on circulating levels of these proteins to effects on LpL, blood pressure, and XXL-VLDL lipoproteins. Supplementary Table S5 shows that *trans*-effects that raise circulating ANGPTL3 raise LpL levels but lower blood pressure and XXL-VLDL lipids. *Trans*-effects that raise circulating ANGPTL4 do the opposite: they lower LpL levels, but raise blood pressure and XXL-VLDL lipids.

## Discussion

Our results identify a set of proteins regulated *in trans* by genetic effects that lower blood pressure and lower triglyceride levels in the XXL-VLDL fraction but raise body mass index: these proteins are encoded by *LPL*, *CD300LG*, *ADIPOQ*, and *TIMP4*. The discordance between the effects on blood pressure and the effects on body mass index indicate that these effects are not mediated by adiposity. The strongest effect of a GATE score on blood pressure was for circulating lipoprotein lipase (LpL). Although circulating LpL is enzymatically inactive ^19^, LpL has been shown to enhance uptake of chylomicrons by hepatocytes by a mechanism that is not dependent on lipolysis ^20^. *CD300LG* is expressed predominantly on endothelial cells where it binds triglyceride-rich lipoproteins ^21^. *ADIPOQ* encodes the secreted protein adiponectin which is taken up by endothelial cells by binding to T-cadherin, where it forms endosomes containing ceramides that are secreted as exosomes ^22^. *TIMP4* knockout in mice alters lipid handling, possibly by post-translational regulation of the fatty acid translocase *CD36* which mediates tissue update of nonesterified fatty acids ^23^.

The associations of blood pressure with genetically predicted levels of lipids in extremely large VLDL (a proxy for chylomicrons) are supported by reported associations with measured levels. In UK Biobank participants, the level of cholesteryl esters in the XXL-VLDL fraction had the strongest association of any metabolite with systolic and diastolic pressure ^9^. In the Women’s Health Study the level of “large VLDL particles” at baseline predicted incident hypertension ^24^.

A likely mechanism by which altered lipid handling might raise blood pressure is through accumulation of lipids in vascular endothelial cells ^25,26^. Endothelial lipid droplet accumulation in response to a high-fat diet reduces nitric oxide production and raises blood pressure ^27^. Accumulation of excess ceramide in endothelial cells also impairs endothelial function ^28^; this may explain the blood pressure-lowering effect of genetic upregulation of adiponectin levels. As accumulation of triglyceride in vascular endothelial cells depends upon lipolysis by membrane-bound LpL, loss-of-function variants in *LPL* would not be expected to increase blood pressure even though they cause hypertriglyceridemia.

The discussion above focuses on proteins regulated by *trans*-QTLs that have discordant effects on blood pressure and on body mass index. A larger group of proteins, including *SHBG* and *F12*, are regulated *in trans* by genetic effects that perturb blood pressure, body mass index and XXL-VLDL-TG in the same direction, it is plausible that these effects on blood pressure also are mediated through pathways related to post-prandial lipid handling. In contrast, proteins including *OGN* and *POMC* are regulated *in trans* by genetic effects that perturb blood pressure and body mass index in the same direction but not XXL-VLDL-TG, indicating that the effects on blood pressure are not mediated through effects on post-prandial lipids. The proteins regulated by *trans*-QTLs that alter blood pressure but not BMI or triglyceride-rich lipoproteins include four that are supported by effects on blood pressure of experimental perturbation in mouse models: *NPPB* (natriuretic peptide B), *VCAN* (versican),

*LGMN* (legumain) and *MFAP4* (microfibrillar-associated protein 4). Of these only *NPPB* is in a pathway targeted by existing antihypertensive drugs. The relation of *LGMN* to blood pressure may be mediated by effects on T cells, as T cell-specific knockout of *LGMN* protects against angiotensin II-induced hypertension ^29^.

### Limitations of this study

The most serious limitation of this study is the incomplete coverage of *trans*-eQTLs in the eQTLGen Phase 1 dataset ^10^. For *trans*-pQTLs we have more complete coverage, but this is limited to the proteins measured on the SomaScan version 4 and Olink Explore 3072 platforms. Until QTL coverage is expanded, it will be difficult to identify which of the many genes influenced by peripheral master regulators such as *SH2B3* mediate effects on blood pressure ^30^. GATE analysis is limited to effects on transcripts in whole blood and proteins in plasma, though many tissue-specific proteins are present in plasma at measurable concentrations. Without access to individual-level proteomics data we cannot directly exclude reverse causation (effect of blood pressure on plasma proteins), which would require re-estimation of SNP-protein summary statistics with restriction to normotensive individuals. However, a causal pathway from genetic regulation of lipid handling to effects on endothelial function and blood pressure is supported by current understanding of mechanisms. Unlike studies based on rare loss-of-function variants, GATE analysis cannot reliably establish the sign of effect of a causal gene, especially where the circulating (soluble) form of the protein may have effects opposite in direction to the effects of signaling through the cellular receptor.

## Perspectives

These results point to a key role in hypertension for genes that regulate post-prandial lipid handling by endothelial cells. The effects on blood pressure of *trans*-QTLs that regulate the levels of ANGPTL4 support a causal role for LpL, as on current understanding the effects of ANGPTL4 are mediated through inhibition of LpL. The genetic evidence for the relationship of blood pressure to lipid handling by endothelial cells is consistent with experimental evidence that lipid accumulation in endothelial cells impairs endothelial function and raises blood pressure. The notion that post-prandial lipid handling has a key role in atherogenesis has a long history ^31^, but there have been few studies of the relation of blood pressure to post-prandial lipid handling. Physical activity lowers both post-prandial lipemia ^32^ and blood pressure ^33^. ANGPTL4 inhibition was reported to lower triglycerides in an early-phase clinical trial, but effects on blood pressure were not reported ^34^.

## Novelty and relevance

### What is new?

- We developed a new genetic method to identify “core” genes on which the small effects of many genetic variants coalesce to influence blood pressure.
- We found a set of proteins where genetic variants that raise levels of the protein in the blood also raise body weight but lower blood pressure and after-meal fat in the blood.

### What is relevant?

- These findings suggest that high blood pressure may be caused by build up of fat in the cells that line blood vessels.

### Clinical/pathophysiological implications?

- One way to control blood pressure is to lower after-meal fat in the blood by exercising more and losing weight. As yet there are no drugs that target this pathway.

## Acknowledgements

This study makes use of de-identified data held by Our Future Health (Study ID OFHS250068). We would like to acknowledge all the research participants who have donated their data to the Our Future Health research programme. Responsibility for interpretation of the data supplied by Our Future Health is the authors’ alone.

## Supplementary Material

### Ethical approval

Ethics approval for Our Future Health was granted by the Cambridge East Research Ethics Committee (REC reference: 21/EE/0016; date of opinion: 29 March 2021). Informed consent was obtained from all participants.

### Calculating QTL-exposure coefficients

The QTL is classified as *cis* or *trans* based on a cutoff of 5 Mb for the distance to the transcription start site of the gene encoding the exposure ((transcript or protein). For each of *K* QTLs for each exposure, the SNP-exposure and SNP-outcome coefficients are adjusted for linkage disequilibrium (LD) by premultiplying by the inverse of the LD matrix between the SNP genotypes in a reference panel (UK Biobank participants of European ancestry).

The QTL-exposure coefficient *α*^*_k_* for the *k*th QTL is the magnitude of the LD-adjusted SNP-exposure coefficient vector. The scalar QTL instrument *Z_k_*is the scalar projection of the genotype vector on the LD-adjusted SNP-exposure coefficient vector.

### Calculating QTL-outcome coefficients and GATE coefficients

The QTL-outcome coefficient *γ*^*_k_* is the ordinary least-squares estimate of the regression of the outcome on *Z_k_*, calculated from the SNP-outcome coefficients from the outcome-GWAS, the LD matrix and allele frequencies of the reference panel. The GATE coefficient and its standard error (SE) for the exposure is calculated from all *K trans*-QTLs as the coefficient of a no-intercept weighted regression of *γ*^*_k_* on *α*^*_k_*, with each QTL weighted by the precision of *γ*^*_k_*. The *cis* coefficient is calculated in the same way but there will usually be only one or *cis*-QTLs.

### Assigning the most likely nearby gene that mediates SNP effects on a trait

The objective of GATE analysis is to identify genes that mediate *in trans* the effects of genetic variants across the genome on a trait, by aggregating *trans*-effects on the expression of that gene as transcript or protein. Although GATE analysis does not depend on being able to identify the nearby genes that mediate *in cis* these *trans*-effects, this identification sometimes provides mechanistic insight. To assign the most likely gene mediating *in cis* the effects of SNPs on traits, we used the Locus-To-Gene model, which scores nearby genes based on proximity of transcription site, variant effect prediction, colocalization of eQTLs, and chromatin interaction features ^16^. For SNP effects on blood pressure, precomputed Locus-To-Gene scores are available on the Open Targets platform for 34 GWAS studies of blood pressure and hypertension. For SNP effects on proteins, we used a reduced version of the Locus-To-Gene model that excluded the colocalization and chromatin-interaction features.

### Supplementary Figures

**Fig S1.**
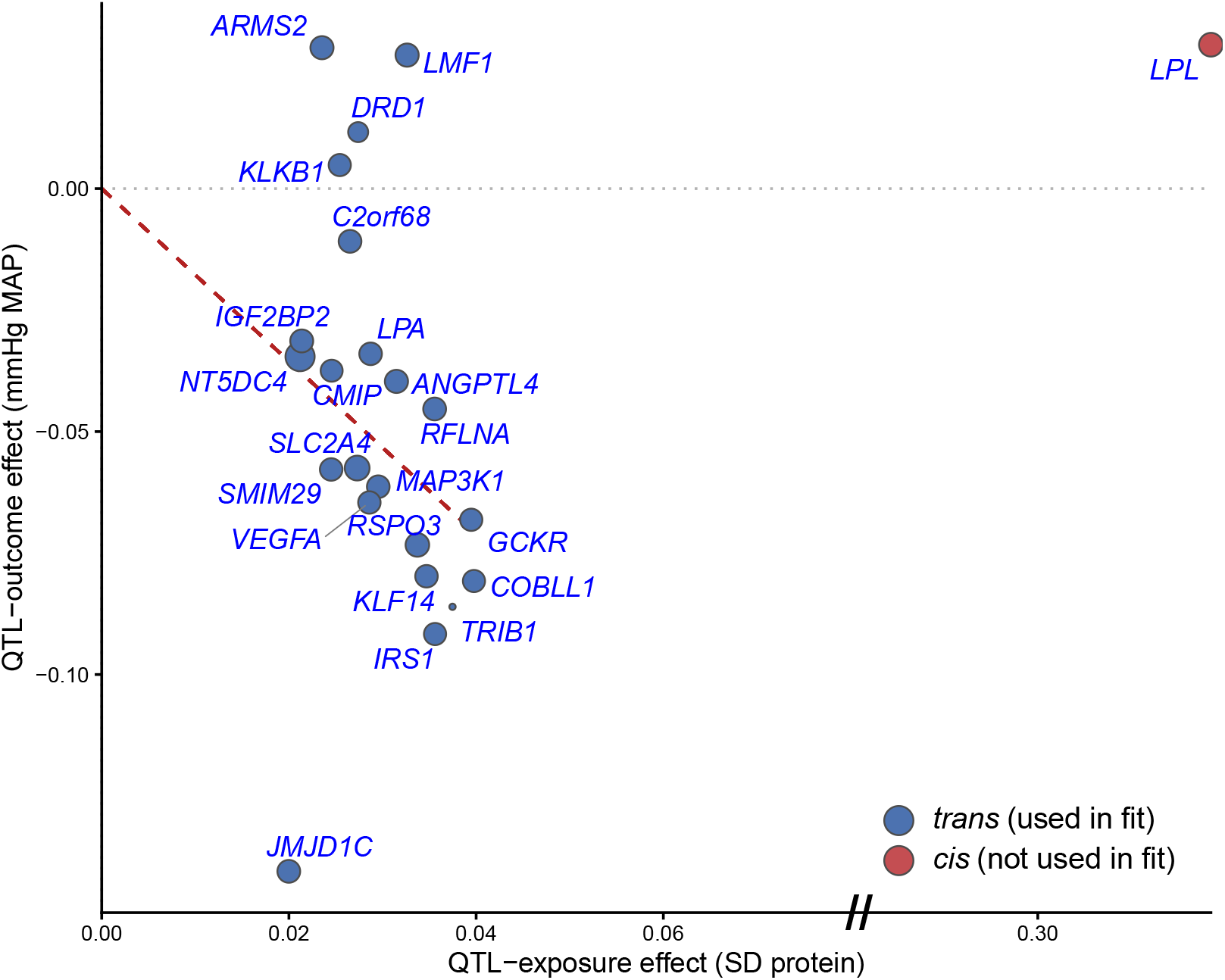
Dose-response scatter plot for the trans- and cis-pQTLs used for *LPL* (Table 2): QTL-exposure effect (SD of circulating LPL) on the x axis against QTL-outcome effect (mmHg MAP) on the y axis, both axes on the same per-SD-of-locus-score scale for every point, cis included. The x axis is broken between 0.08 and 0.30 (marked ”//”), a gap containing no data, so that the 22 *trans*-pQTLs (all below 0.04) are not compressed against the origin by the single outlying *cis*-pQTL (≈ 0.32). Point size is inversely proportional to the standard error of the QTL-exposure effect. The *cis*-pQTL (red) is shown for comparison only and was not used to fit the model. The slope of the fitted line (dashed red) is the maximum likelihood estimate of the causal effect parameter, drawn only across the trans loci actually fitted (not extended to the *cis* point). Labels (positioned to avoid overlap) show the most likely gene at each locus, assigned by reduced Locus-To-Gene procedure.

**Fig S2.**
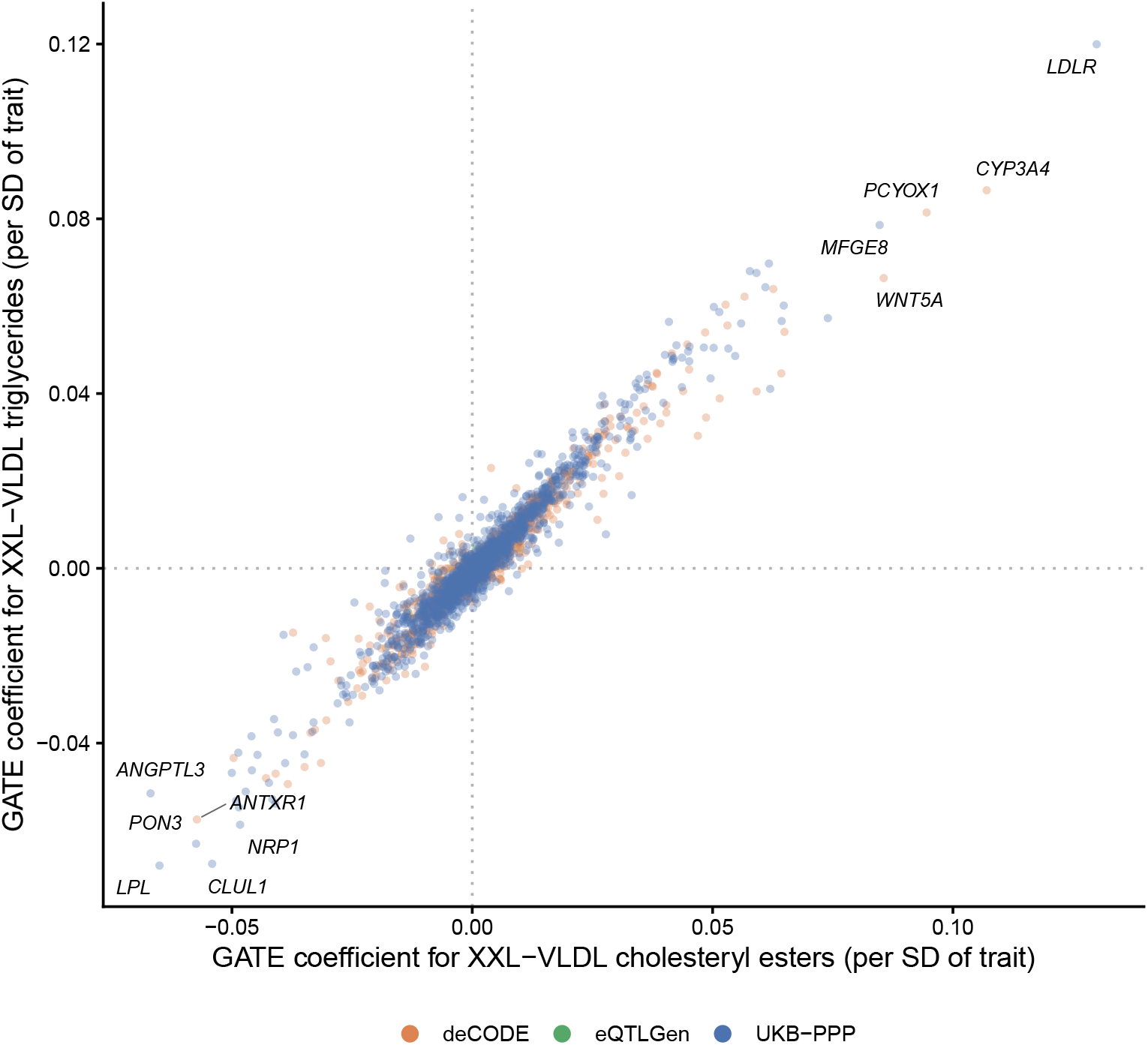
GATE coefficients for XXL-VLDL cholesteryl esters (x axis) against XXL-VLDL triglycerides (y axis; Karjalainen et al. 2024, GWAS Catalog GCST90302164 and GCST90302172 respectively, same 33-cohort NMR-metabolomics study), for the 4000 exposures with *N*_eff_ *>* 5 in both screens (Pearson *r* = 0.965), with the chr 11 APOA5 locus excluded from both. Points are colored by the transcriptomic/proteomic study (eQTLGen, UKB-PPP or deCODE) from which the exposure was drawn; the 11 exposures at the extremes of either axis are labeled.

### Supplementary Tables

**Table S1.** Leave-one-locus-out (LOLO) sensitivity analysis for the 27 exposures in Table 2 with *p*_GATE_ *<* 1 × 10^−6^.

| Protein | $K$ | $\hat{\beta}_{\text{GATE}}$ | LOLO range | Most influential locus | | | | |
| --- | --- | --- | --- | --- | --- | --- | --- | --- |
| | | | | Chr | Position (Mb) | Likely gene | $\hat{\beta}_{\text{LOLO}}$ | $p_{\text{LOLO}}$ |
| *POMC* | 26 | -0.163 | [-0.188, -0.112] | 16 | 81.41 | *CMIP* | -0.112 | $3 \times 10^{-10}$ |
| *CCL24* | 33 | 0.154 | [0.049, 0.169] | 11 | 45.85 | *SPI1* | 0.049 | 0.006 |
| *NPPB* | 17 | -0.146 | [-0.169, -0.113] | 10 | 73.65 | *SYNPO2L* | -0.113 | $2 \times 10^{-10}$ |
| *SPOCK1* | 28 | -0.132 | [-0.144, -0.113] | 11 | 16.21 | *SOX6* | -0.113 | $3 \times 10^{-10}$ |
| *VCAN* | 40 | -0.147 | [-0.160, -0.085] | 4 | 98.71 | *SLC39A8* | -0.085 | $2 \times 10^{-6}$ |
| *ADGRG2* | 25 | -0.140 | [-0.152, -0.109] | 4 | 101.78 | *SLC39A8* | -0.109 | $1 \times 10^{-9}$ |
| *C9* | 17 | -0.132 | [-0.155, -0.105] | 5 | 56.50 | *MAP3K1* | -0.105 | $5 \times 10^{-9}$ |
| *ADIPOQ* | 23 | -0.153 | [-0.170, -0.113] | 1 | 11.78 | *MTHFR* | -0.113 | $2 \times 10^{-10}$ |
| *DSG2* | 32 | -0.138 | [-0.156, -0.103] | 4 | 101.63 | *BANK1* | -0.103 | $8 \times 10^{-9}$ |
| *CD300LG* | 43 | -0.137 | [-0.153, -0.107] | 4 | 101.63 | *SLC39A8* | -0.107 | $2 \times 10^{-9}$ |
| *LPL* | 22 | -0.223 | [-0.236, -0.206] | 10 | 63.14 | *JMJD1C* | -0.206 | $1 \times 10^{-30}$ |
| *TIMP4* | 22 | -0.129 | [-0.147, -0.110] | 2 | 226.00 | *IRS1* | -0.110 | $7 \times 10^{-10}$ |
| *IL17RB* | 34 | -0.166 | [-0.172, -0.140] | 14 | 94.03 | *SERPINA1* | -0.140 | $5 \times 10^{-15}$ |
| *CKB* | 22 | -0.133 | [-0.165, -0.112] | 4 | 3.39 | *HGFAC* | -0.165 | $3 \times 10^{-20}$ |
| *SHBG* (deCODE) | 19 | -0.151 | [-0.167, -0.101] | 17 | 49.17 | *ZNF652* | -0.101 | $2 \times 10^{-8}$ |
| *SHBG* (UKB-PPP) | 31 | -0.141 | [-0.162, -0.100] | 10 | 62.16 | *JMJD1C* | -0.100 | $2 \times 10^{-8}$ |
| *HTRA1* | 14 | 0.139 | [0.097, 0.164] | 4 | 99.14 | *ADH4* | 0.097 | $5 \times 10^{-8}$ |
| *NAGPA* | 19 | 0.132 | [0.110, 0.147] | 10 | 63.12 | *JMJD1C* | 0.110 | $8 \times 10^{-10}$ |
| *RNASE4* | 14 | 0.143 | [0.115, 0.165] | 4 | 3.29 | *HGFAC* | 0.115 | $1 \times 10^{-10}$ |
| *CNTN3* | 51 | 0.156 | [0.135, 0.170] | 11 | 16.21 | *SOX6* | 0.135 | $4 \times 10^{-14}$ |
| *IGSF9* | 16 | 0.133 | [0.114, 0.145] | 2 | 210.67 | *CPS1* | 0.114 | $2 \times 10^{-10}$ |
| *F12* | 20 | 0.144 | [0.109, 0.152] | 10 | 63.06 | *REEP3* | 0.109 | $1 \times 10^{-9}$ |
| *SERPINA4* | 15 | 0.128 | [0.095, 0.139] | 4 | 3.25 | *DOK7* | 0.095 | $1 \times 10^{-7}$ |
| *EDAR* | 20 | -0.130 | [-0.140, -0.083] | 10 | 100.98 | *SUFU* | -0.083 | $3 \times 10^{-6}$ |
| *EFCAB14* | 13 | 0.167 | [0.080, 0.189] | 15 | 90.85 | *FURIN* | 0.080 | $7 \times 10^{-6}$ |
| *OGN* | 24 | 0.154 | [0.134, 0.174] | 20 | 41.00 | *CHD6* | 0.174 | $2 \times 10^{-22}$ |
| *VWC2* | 20 | 0.139 | [0.120, 0.147] | 7 | 151.71 | *PRKAG2* | 0.120 | $2 \times 10^{-11}$ |

$K$ : number of trans-pQTLs. $\hat{\beta}_{\text{GATE}}$ : GATE coefficient; LOLO range: [min, max] of the $K$ GATE estimates obtained by omitting one locus at a time. Most influential locus: the locus for which omission changes $\hat{\beta}_{\text{GATE}}$ the most. Chr and Position (Mb): chromosome and base position of the locus window start. Likely gene assigned by reduced Locus-To-Gene model. $\hat{\beta}_{\text{LOLO}}$ and $p_{\text{LOLO}}$ : GATE estimate and $p$ -value when that locus is omitted.

**Table S2.** Replication, in an independent GWAS of systolic blood pressure with no overlap with UK Biobank (GERA cohort), of the GATE score associations with mean arterial pressure for the 27 exposures in Table 2.

| Protein | MAP (OFH, discovery)<br>N = 373,882 |  | SBP (GERA, replication)<br>N = 99,785 |  |
| --- | --- | --- | --- | --- |
|  | Estimate | *p* | Estimate | *p* |
| *LPL* | -0.223 | $9 \times 10^{-36}$ | -0.234 | $9 \times 10^{-9}$ |
| *EFCAB14* | 0.167 | $8 \times 10^{-21}$ | 0.098 | 0.02 |
| *IL17RB* | -0.166 | $2 \times 10^{-20}$ | -0.133 | 0.001 |
| *POMC* | -0.163 | $9 \times 10^{-20}$ | -0.159 | $1 \times 10^{-4}$ |
| *CNTN3* | 0.156 | $3 \times 10^{-18}$ | 0.183 | $7 \times 10^{-6}$ |
| *CCL24* | 0.154 | $7 \times 10^{-18}$ | 0.081 | 0.05 |
| *OGN* | 0.154 | $9 \times 10^{-18}$ | 0.038 | 0.3 |
| *ADIPOQ* | -0.153 | $1 \times 10^{-17}$ | -0.176 | $2 \times 10^{-5}$ |
| *SHBG* (deCODE) | -0.151 | $3 \times 10^{-17}$ | -0.119 | 0.003 |
| *VCAN* | -0.147 | $2 \times 10^{-16}$ | -0.097 | 0.02 |
| *NPPB* | -0.146 | $3 \times 10^{-16}$ | -0.198 | $1 \times 10^{-6}$ |
| *F12* | 0.144 | $7 \times 10^{-16}$ | 0.073 | 0.07 |
| *RNASE4* | 0.143 | $1 \times 10^{-15}$ | 0.066 | 0.1 |
| *SHBG* (UKB-PPP) | -0.141 | $3 \times 10^{-15}$ | -0.091 | 0.03 |
| *ADGRG2* | -0.140 | $5 \times 10^{-15}$ | -0.075 | 0.07 |
| *VWC2* | 0.139 | $7 \times 10^{-15}$ | 0.112 | 0.006 |
| *HTRA1* | 0.139 | $7 \times 10^{-15}$ | 0.125 | 0.002 |
| *DSG2* | -0.138 | $1 \times 10^{-14}$ | -0.129 | 0.002 |
| *CD300LG* | -0.137 | $2 \times 10^{-14}$ | -0.137 | $8 \times 10^{-4}$ |
| *IGSF9* | 0.133 | $1 \times 10^{-13}$ | 0.152 | $2 \times 10^{-4}$ |
| *CKB* | -0.133 | $1 \times 10^{-13}$ | -0.098 | 0.02 |
| *C9* | -0.132 | $1 \times 10^{-13}$ | -0.096 | 0.02 |
| *NAGPA* | 0.132 | $1 \times 10^{-13}$ | 0.130 | 0.001 |
| *SPOCK1* | -0.132 | $2 \times 10^{-13}$ | -0.016 | 0.7 |
| *EDAR* | -0.130 | $3 \times 10^{-13}$ | -0.036 | 0.4 |
| *TIMP4* | -0.129 | $6 \times 10^{-13}$ | -0.169 | $3 \times 10^{-5}$ |
| *SERPINA4* | 0.128 | $8 \times 10^{-13}$ | 0.100 | 0.01 |

GERA: Hoffmann et al. 2017, GWAS Catalog accession GCST007095; N=99,785; long-term-average systolic blood pressure (mean of repeated clinic readings), raw mmHg scale. Both the MAP (OFH) and SBP (GERA) GATE scores exclude *trans*-QTLs in the HLA, *SH2B3* and 8p23.1 inversion regions. GATE coefficients are the change in the outcome (mmHg) for an increase of the GATE score by 1 SD.

**Table S3.** Trans- and cis-pQTLs for *LPL* (Table 2), underlying the dose-response analysis shown in Figure S1.

| Genomic region |  | SNPs | Locus type | Effect on MAP | *p* | Likely gene | Full gene name |
| --- | --- | --- | --- | --- | --- | --- | --- |
| Chr | Mb (GRCh38) |  |  |  |  |  |  |
| 2 | 27.15-27.61 | 28 | trans | -0.068 | $1 \times 10^{-4}$ | *GCKR* | glucokinase regulator |
| 2 | 85.56-85.65 | 11 | trans | -0.011 | 0.5 | *C2orf68* | chromosome 2 open reading frame 68 |
| 2 | 112.73-112.73 | 1 | trans | -0.035 | 0.05 | *NT5DC4* | 5'-nucleotidase domain containing 4 |
| 2 | 164.65-164.88 | 62 | trans | -0.081 | $6 \times 10^{-6}$ | *COBLL1* | cordon-bleu WH2 repeat protein like 1 |
| 2 | 226.00-226.33 | 117 | trans | -0.092 | $3 \times 10^{-7}$ | *IRS1* | insulin receptor substrate 1 |
| 3 | 185.78-185.82 | 40 | trans | -0.031 | 0.08 | *IGF2BP2* | insulin like growth factor 2 mRNA binding protein 2 |
| 4 | 186.22-186.26 | 27 | trans | 0.005 | 0.8 | *KLKB1* | kallikrein B1 |
| 5 | 56.56-56.57 | 11 | trans | -0.061 | $6 \times 10^{-4}$ | *MAP3K1* | mitogen-activated protein kinase kinase kinase 1 |
| 5 | 175.22-175.22 | 3 | trans | 0.012 | 0.5 | *DRD1* | dopamine receptor D1 |
| 6 | 34.13-34.39 | 50 | trans | -0.058 | 0.001 | *SMIM29* | small integral membrane protein 29 |
| 6 | 43.79-43.80 | 5 | trans | -0.065 | $3 \times 10^{-4}$ | *VEGFA* | vascular endothelial growth factor A |
| 6 | 126.49-127.21 | 8 | trans | -0.073 | $4 \times 10^{-5}$ | *RSPO3* | R-spondin 3 |
| 6 | 160.26-160.66 | 96 | trans | -0.034 | 0.06 | *LPA* | lipoprotein(a) |
| 7 | 130.74-130.78 | 64 | trans | -0.080 | $8 \times 10^{-6}$ | *KLF14* | KLF transcription factor 14 |
| 8 | 19.29-21.22 | 920 | cis | 0.030 | 0.1 | *LPL* | lipoprotein lipase |
| 8 | 125.46-125.50 | 48 | trans | -0.086 | $2 \times 10^{-6}$ | *TRIB1* | tribbles pseudokinase 1 |
| 10 | 63.14-63.59 | 76 | trans | -0.141 | $4 \times 10^{-15}$ | *JMJD1C* | jumonji domain containing 1C |
| 10 | 122.45-122.45 | 1 | trans | 0.029 | 0.1 | *ARMS2* | age-related maculopathy susceptibility 2 |
| 12 | 123.14-124.13 | 54 | trans | -0.045 | 0.01 | *RFLNA* | refilin A |
| 16 | 0.90-0.99 | 139 | trans | 0.027 | 0.1 | *LMF1* | lipase maturation factor 1 |
| 16 | 81.49-81.50 | 3 | trans | -0.037 | 0.04 | *CMIP* | c-Maf inducing protein |
| 17 | 7.23-7.30 | 7 | trans | -0.058 | 0.001 | *SLC2A4* | solute carrier family 2 member 4 |
| 19 | 7.90-8.55 | 4 | trans | -0.040 | 0.03 | *ANGPTL4* | angiopoietin like 4 |

**Table S4.**
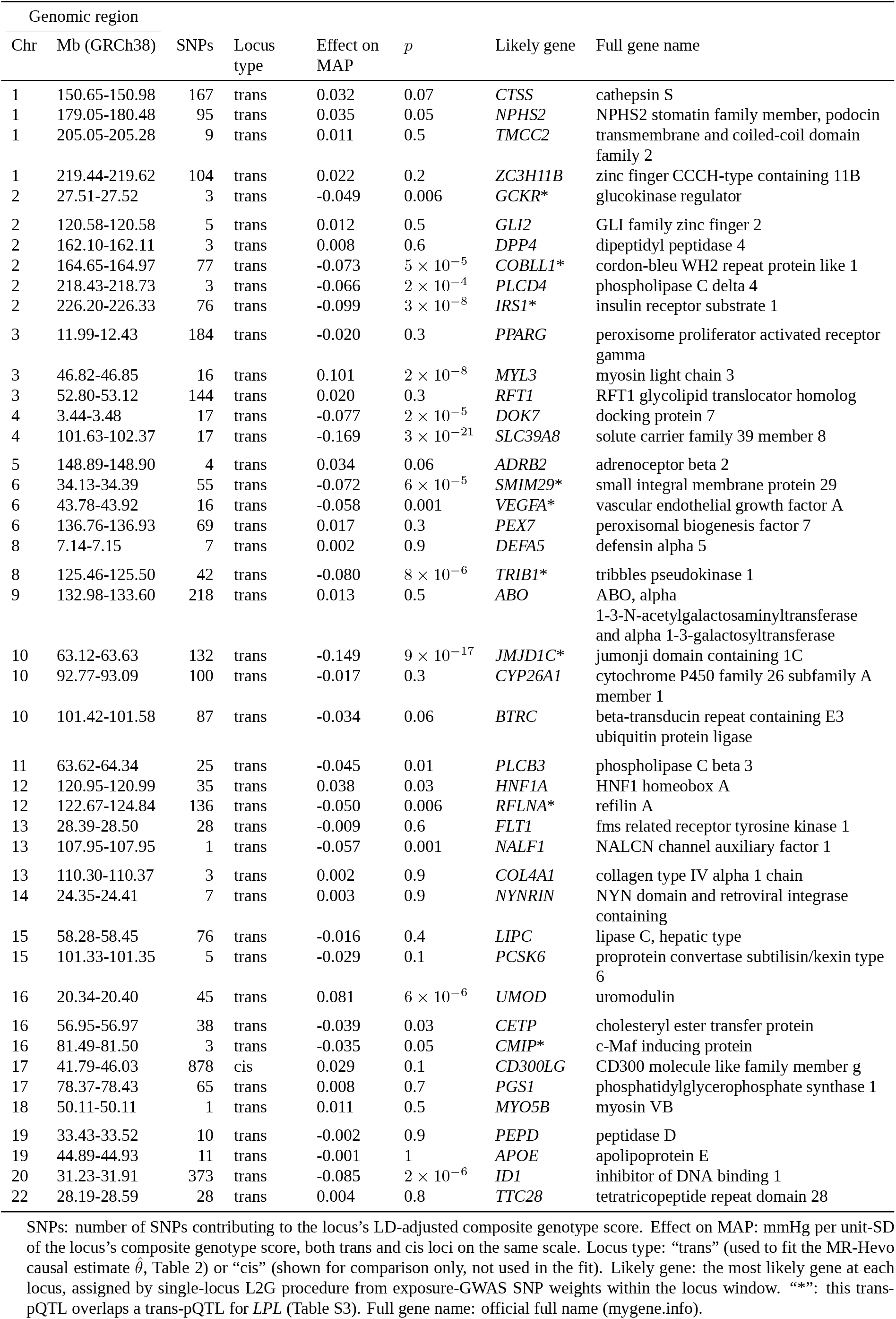
Trans- and cis-pQTLs for *CD300LG* (Table 2).

**Table S5.**
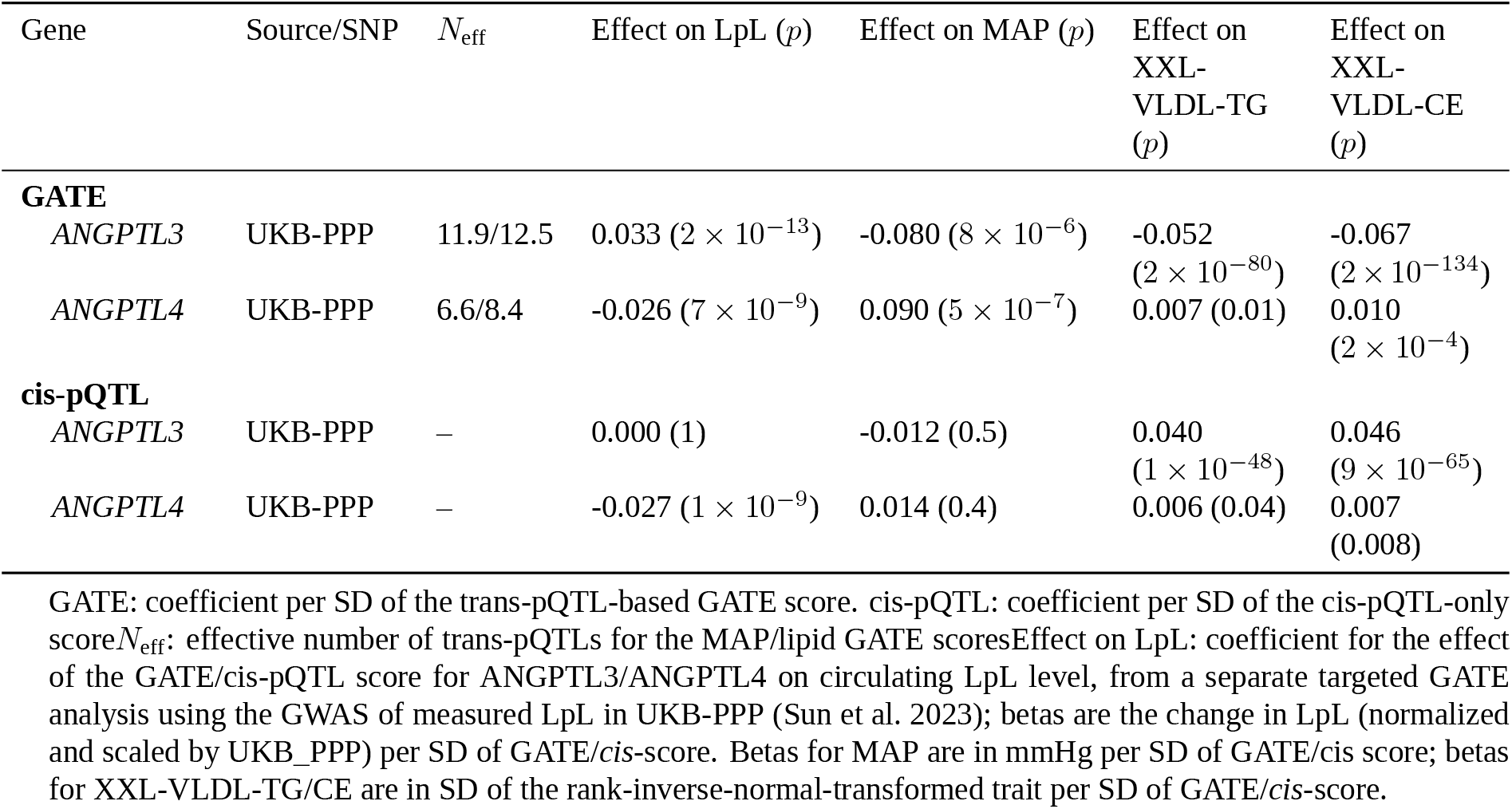
Genetic effects on *ANGPTL3* and *ANGPTL4* – post-translational regulators of lipoprotein lipase – on circulating LpL, mean arterial pressure, and XXL-VLDL lipid traits, combining GATE score and cis-pQTL-score.

| Gene | Source/SNP | $N_{\text{eff}}$ | Effect on LpL ($p$) | Effect on MAP ($p$) | Effect on<br>XXL-VLDL-TG<br>($p$) | Effect on<br>XXL-VLDL-CE<br>($p$) |
| --- | --- | --- | --- | --- | --- | --- |
| **GATE** |  |  |  |  |  |  |
| *ANGPTL3* | UKB-PPP | 11.9/12.5 | 0.033 ($2 \times 10^{-13}$) | -0.080 ($8 \times 10^{-6}$) | -0.052<br>($2 \times 10^{-80}$) | -0.067<br>($2 \times 10^{-134}$) |
| *ANGPTL4* | UKB-PPP | 6.6/8.4 | -0.026 ($7 \times 10^{-9}$) | 0.090 ($5 \times 10^{-7}$) | 0.007 (0.01) | 0.010<br>($2 \times 10^{-4}$) |
| **cis-pQTL** |  |  |  |  |  |  |
| *ANGPTL3* | UKB-PPP | – | 0.000 (1) | -0.012 (0.5) | 0.040<br>($1 \times 10^{-48}$) | 0.046<br>($9 \times 10^{-65}$) |
| *ANGPTL4* | UKB-PPP | – | -0.027 ($1 \times 10^{-9}$) | 0.014 (0.4) | 0.006 (0.04) | 0.007<br>(0.008) |

GATE: coefficient per SD of the trans-pQTL-based GATE score. cis-pQTL: coefficient per SD of the cis-pQTL-only score $N_{\text{eff}}$ : effective number of trans-pQTLs for the MAP/lipid GATE scores Effect on LpL: coefficient for the effect of the GATE/cis-pQTL score for *ANGPTL3*/*ANGPTL4* on circulating LpL level, from a separate targeted GATE analysis using the GWAS of measured LpL in UKB-PPP (Sun et al. 2023); betas are the change in LpL (normalized and scaled by UKB\_PPP) per SD of GATE/*cis*-score. Betas for MAP are in mmHg per SD of GATE/*cis* score; betas for XXL-VLDL-TG/CE are in SD of the rank-inverse-normal-transformed trait per SD of GATE/*cis*-score.

**Table S6.**
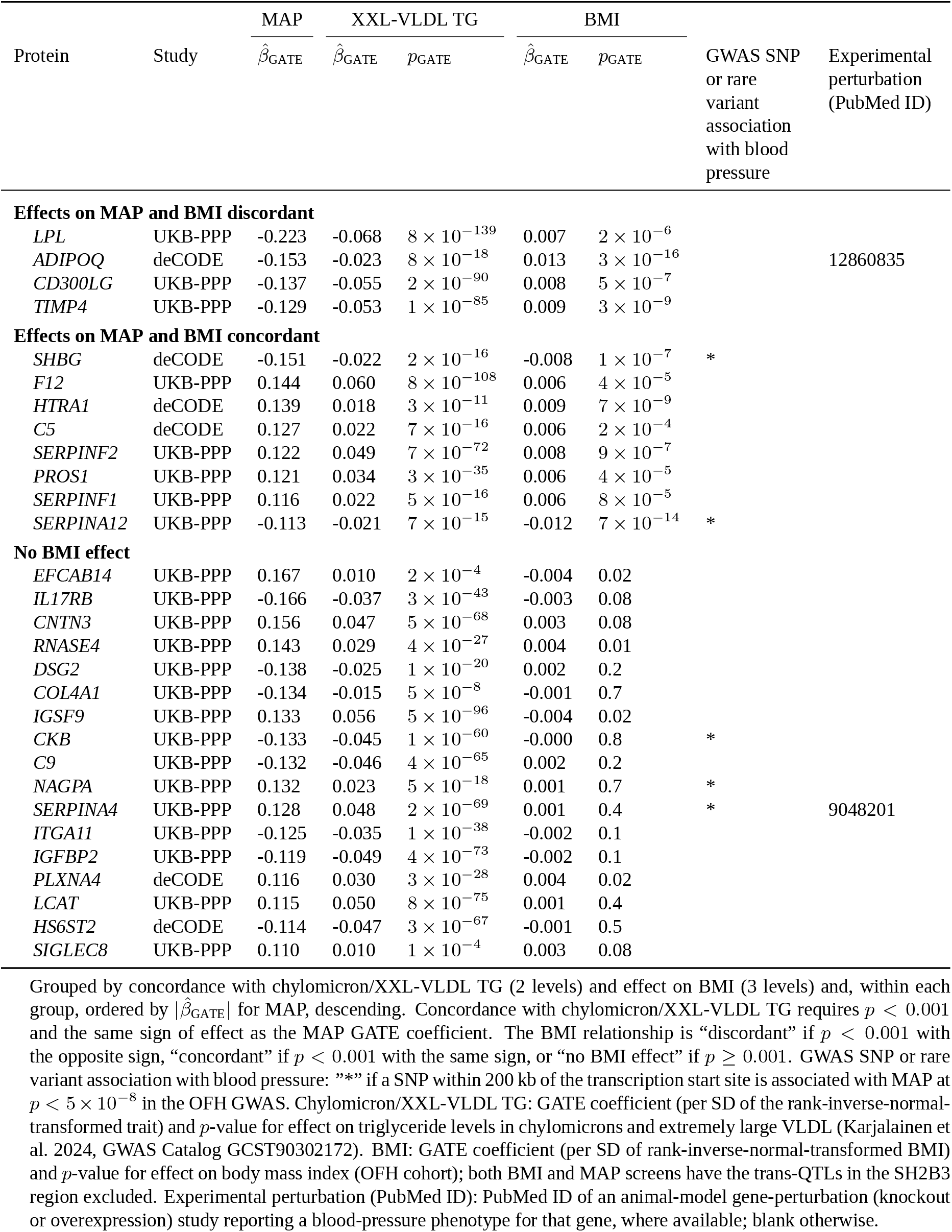
Effect on XXL-VLDL triglycerides and on body mass index for the 29 genes with *N*_eff_ *>* 5 and GATE *p* < 10^−9^ for GATE association with MAP, surviving a leave-one-locus-out (LOLO) sensitivity analysis at *p* < 10^−5^ after omitting the single most influential trans QTL and with effect on XXL-VLDL-TG concordant with effect on MAP.

**Table S7.**
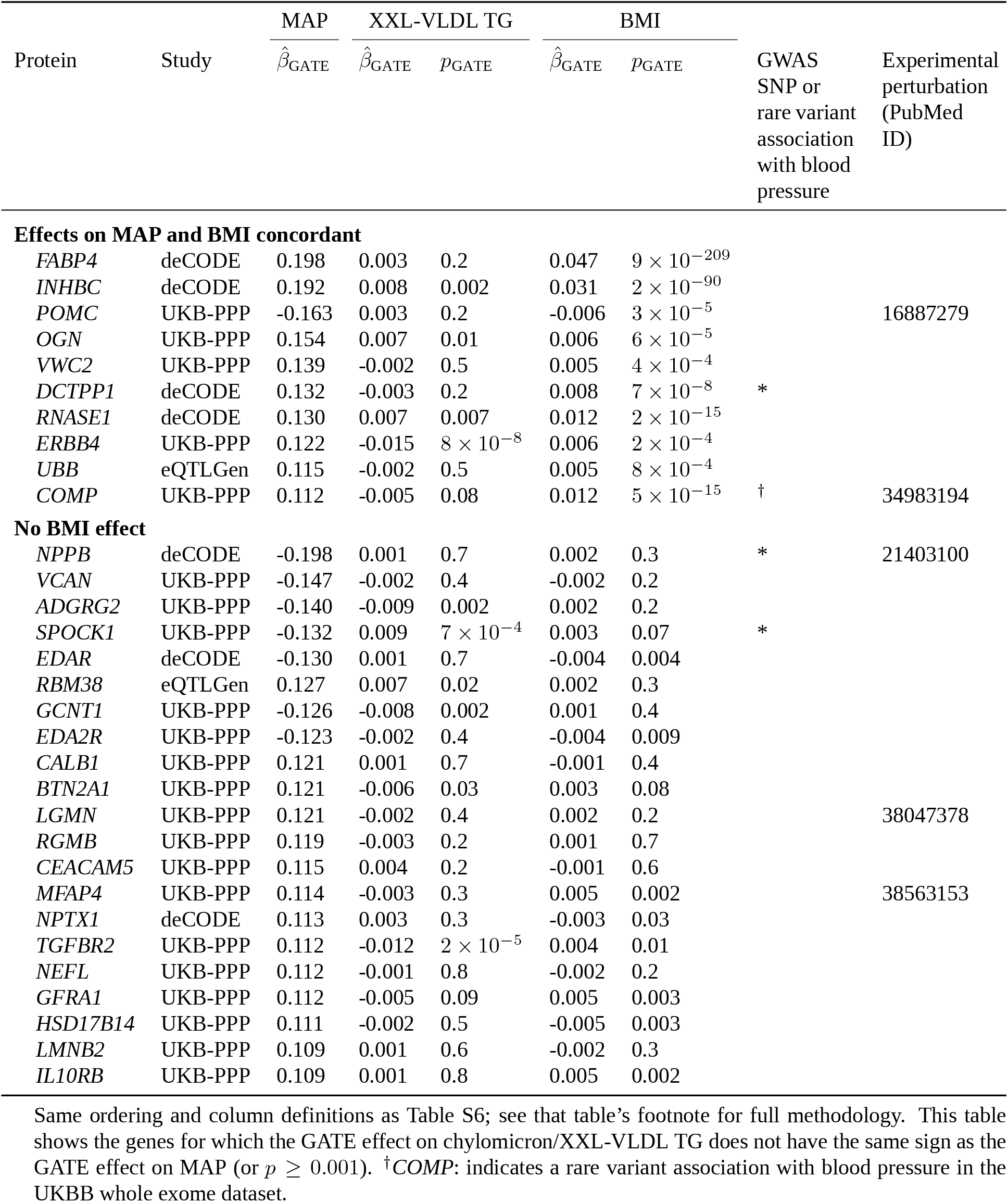
Effect on XXL-VLDL triglycerides and on body mass index for the 31 genes with *N*_eff_ *>* 5 and GATE *p* < 10^−9^ for GATE association with MAP, surviving a leave-one-locus-out (LOLO) sensitivity analysis at *p* < 10^−5^ after omitting the single most influential trans QTL and without effect on XXL-VLDL-TG concordant with effect on MAP.

